# Is water vole diet consistent with the *plant hypothesis* for explaining population fluctuations?

**DOI:** 10.1101/2024.09.04.611276

**Authors:** Hélène Lisse, Marion Buronfosse, Cédric Jacquet, Gaëlle Sobczyk-Moran, Etienne Ramadier, Ambre Fafournoux, Virginie Lattard, Adrien Pinot

**Author notes:** Corresponding author: Adrien Pinot.

## Abstract

Rodent population cycles are observed in highly seasonal environments. As most rodents are herbivorous, the availability and the quality of their food resources varies greatly across seasons. Furthermore, it is well documented that herbivore densities have a measurable effect on vegetation and conversely. So, many studies investigated whether rodent population cycles could be induced by bottom-up regulation. A recent review summarized several sub-hypotheses leading to rodent population cycles: cycles may be due to inherent inter-annual variations of plant quantity, to overshoot of carrying capacity by overgrazing (i.e. lack of quantity), to changes in quality of food (decrease of quality of preferred food or switch towards less quality food) in response to rodent grazing (e.g. plant defences). If some sub-hypothesis seems to be more important than others, there is currently a prerequisite to construct scientific consensus: dietary description is still overlooked in many systems and should be more investigated.

This study focuses on fossorial water vole. It shows contrasted population dynamics depending on its geographical locations. It is known to be able to exhibit large outbreaks in grasslands in highly seasonal climate. It is thus a good model species to investigate plant hypotheses, first beginning by diet description.

The diet of water vole was investigated in and out of the outbreak area with a combination of approaches in the field, in different sampling sites and considering seasonality. We demonstrated that voles have a very large fundamental trophic niche, but strong behavioural selection, inducing a narrower realised niche, especially during winter. We created an experimental device based on camera trap and cafeteria tests. We observed a strong preference for dandelion (*Taraxacum officinale*) in wild water voles, that results in exclusive selection during winter for food stores. These preferences were constant across seasons, altitudes and grassland productivity gradients, despite the scarcity of this species in some experimental sites.

First, we conclude on the importance of using different methods to fully describe the diet of rodents Second, we assess that dandelion is a winter key resource for water vole. It thus might be interesting to investigate the role of dandelion in vole population dynamics.

## Introduction

Rodents are an important model for studying population dynamics. Numerous studies have explored the mechanisms underlying fluctuations in their numbers, particularly in the context of cyclical dynamics. Several reference books have been regularly published on the subject (e.g. *Elton 1942*, *Royama 1992*, *Berryman 2002*, *Turchin 2003*, *Krebs 2013*), but it is clear that further work is needed to fully understand the relative importance of the different hypotheses in the general pattern of cycles (*Oli 2019*, *Andreassen et al. 2021*, *Soininen and Neby 2023*). There is, however, a scientific consensus that trophic interactions are important drivers of cycles (*Berryman 2002*), that cycles occur in seasonal environment (*Kendal et al. 1998*, *Tkadlec and Stenseth 2001*), and that delayed density dependence is necessary to observe long-phase cycles (*Hornfeldt 1994*, *Stenseth 1999*).

Seasonal environments are typically characterised by simplified trophic chains. This can result in the establishment of prey-specialist predator relationships, favored by the presence of snow cover (*Hansson and Henttonen 1985*). The delay in this process is induced by the generation time of specialist predators (*Hanski et al. 1991*). Seasonality also implies significant variation in food resource throughout the year. Rodent-plant interactions may also be highly detrimental to plants during winter, when they cannot regrow, yet rodents still need to feed. As a result, rodents can significantly impact the quantity (e.g., *Huitu et al. 2003*) or quality (through the synthesis of secondary metabolites, e.g. *Massey et al. 2007*) of their resources. The time required for the regeneration of preferred resources after depletion, or for the restoration of food quality following plant defenses, can theoretically induce cycles (e.g *Turchin et al. 2000*, *Massey et al. 2008*, for a review, see also *Soininen and Neby 2023*). In both cases, rodents must depend on a preferred food and the absence of this resource must impact their survival, reproduction or dispersal (*Soininen and Neby 2023*). Plant-rodent interactions have been proposed to explain certain rodent cycles in several different contexts (e. g., *Lemmus* – mosses: *Turchin et al. 2000*, *Lemmus* – carex: *Seldal et al. 1994*, *Microtus* – graminoids: *Massey et al. 2008*, *Myodes* – blueberry: *Selås 2020*). However, scientific consensus has not yet been completely achieved in these systems. One explanation is that studying population dynamics in the wild requires complex data that are often difficult to access. Further research is needed to determine the extent to which observed cycles are driven by plant-rodent relationships.

To achieve a cycle driven by plant-rodent interactions, candidate systems must meet the 3 following points: i) the plants exhibit strong seasonality in their growth period, ii) the rodents have strong food preference or are dependent on a limited number of plant species, iii) after the rodents have affected their preferred food source, a recovery time is required to restore its quality or quantity.

In the French mid-mountains (Jura and Massif Central), water voles exhibit some prerequisites for plant-based hypotheses. At first, outbreaks occur only above 750 meters above the sea (*Fichet-Calvet et al. 2000*) in areas where there is significant seasonality in grassland growth. Furthermore, there is a wide diversity of grasslands in the Massif Central mountains, due to gradients in altitude, productivity, soil hygrometry and type of harvesting (for a full description, see *Galliot et al. 2020*). Depending on these gradients, the flora may change completely from one agricultural plot to another. Field observations suggest that grasslands sensitivity to outbreaks varies among different types of grassland. Indeed, low-productive grasslands seem to support lower vole densities during outbreaks *(Fichet-Calvet et al. 2000*). Moreover, outbreaks significantly change plant communities (*Nicod et al. 2020*) highlighting a close link between plant and water vole population.

However, our understanding of the water vole’s diet remains limited, mainly due to the challenges of studying the feeding habits of subterranean animals within a complex environment. While it is scientifically agreed that the water vole is strictly herbivorous, its precise diet is not clearly described. Airoldi (1976) and Potapov *et al.* (2004) found voles’ winter food stores. It was composed of roots from various species in the former study. The specific contents were not described in the latter. In both cases, the weight of these food stores was consequent reaching hundreds of grams or kilograms and are season dependent (i.e. discovered in fall and winter). Several other studies have indicated that water voles exhibit food preferences (*Viette et al. 2000*; *Saucy et al. 1999*; *Koop 1988*). Viette *et al.* (2000) and Saucy *et al.* (1999) found that these preferences occur at the intraspecific scale in response to variations in plant chemical traits, particularly plant defenses. Koop (1988) observed that food preferences among grassland species influenced the voles’ foraging behavior and space use, with a notable preference for dandelions and clovers over poaceae. However, while the study design was relevant, its conclusions were constrained by the limited number of plant species considered and the lack of replication. A more recent study did a large investigation of water vole diet in wild; it revealed that voles have a diverse diet with Poaceae being dominant, exhibit strong site-specific variations and exhibit a slight seasonal pattern (Øtsby 2019). Furthermore, Neyland (2011) showed that water voles make food choices likely linked to the chemical traits of plants, particularly their nitrogen content. All these studies suggest that the water vole’s diet results from a complex interplay of factors, including habitat and seasonal variations. It also shows that water vole is sensitive to food chemical traits and make choices.

In this study, the trophic niche was investigated as part of the ecological niche including potentiality of vole’s diet and behavioural preferences for some aliments, in natural environment. We used definitions from the theoretical framework described by Shipley and collaborators (*2009*, see fig 1). The term **fundamental trophic niche** hereafter in this paper describes voles’ eating possibilities regardless of habitat or season. In contrast, the **realised trophic niche** defines the part of the fundamental trophic niche that is determined by habitat and feeding behaviour under natural living conditions. The realised niche obviously depends on the **available niche** (plants that the vole encounters in its habitat) and on the vole’s **preferences** (plants that the vole chooses to eat).

**Figure 1:**
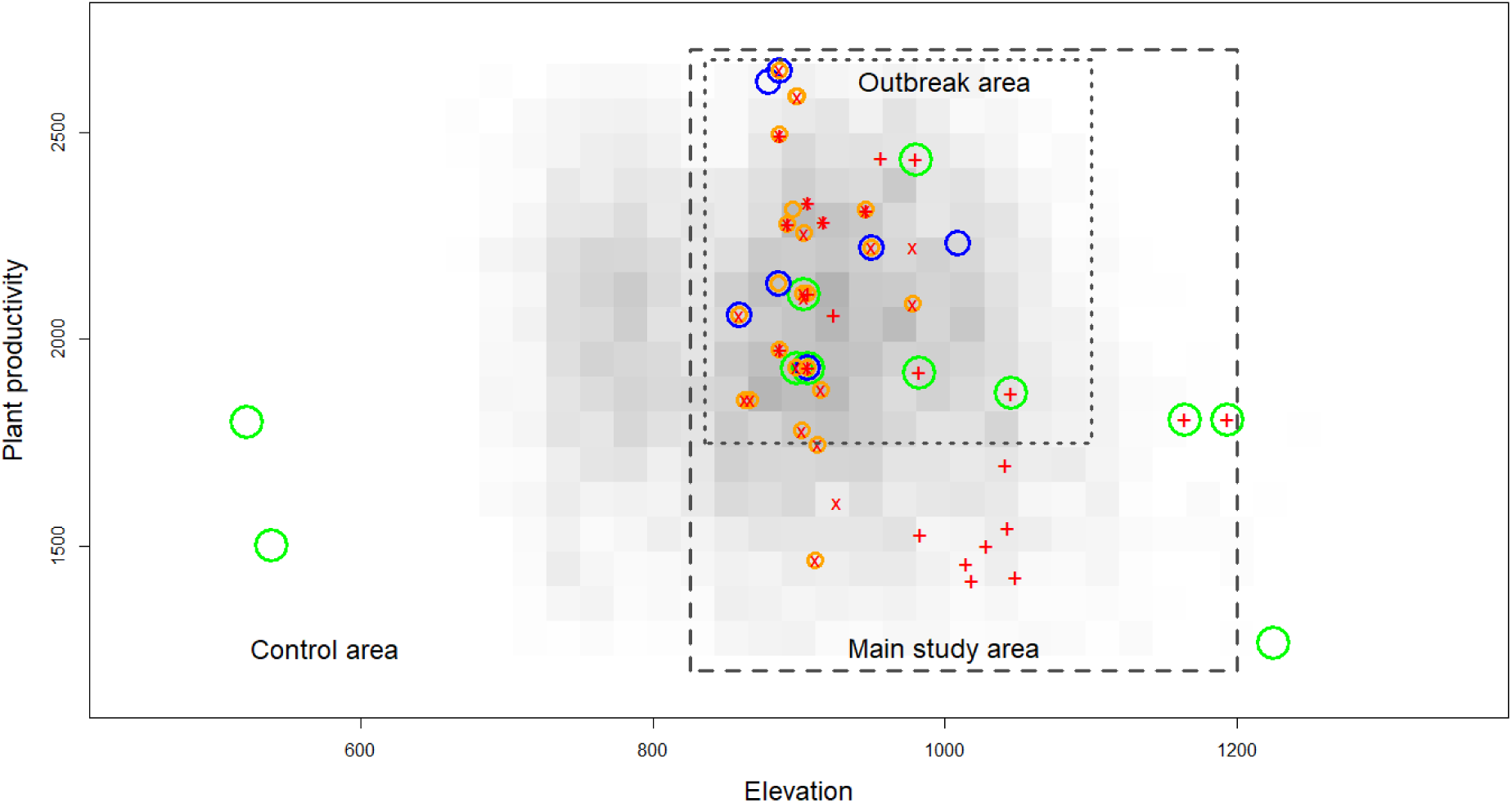
Localisation of sampled plots in space (A) and on plant productivity (sentinel data summarised at field plot scale) and altitudinal gradient (B). Crosses represent locations of environmental data samplings for plant/root availability. Rings represent locations of vole’s samplings. Food choices (green rings) were studied with the Campascope in field. Food stores (blue rings) were found empirically during vole trapping. Stomach contents (orange rings) were collected during trapping sessions. The number in brackets in the legend corresponds to the number of plots (which may be smaller than the number of samples, since in some cases several samples were taken on the same plot). Grey squares represent the density of field plot in our main study area and in its margins (the more grey is the most abundant, total number of field plot : 9000).

Formerly, many designs have been explored in dietary studies to assess the trophic niche of rodents. Authors usually used direct observations of vole stomachs or faeces contents with species determination under the microscope (Hansson 1970), such as Cole and Batzli (1979) on prairie vole, Ashby and Vincent (1976) on the aquatic form of water vole, Pinter *et al.* (1992) on Montane vole and Rothstein and Tamarin (1977) on insular beach vole. More recently, DNA amplification and sequencing for species determination in stomachs or faecal contents have been used for comparison with DNA bases *(Taberlet et al. 2012*), giving good results for dietary study in small herbivores and rodents (*Soininen et al. 2009*), such as for the fossorial form of water vole in Norway (*Østby 2019*), Amargosa vole in California (*Castle et al, 2020*), Norwegian lemming in Norway (*Soininen et al. 2017*) or bank and tundra vole in Norway (*Neby et al. 2024)*. Moreover, voles are known to have consistent food stores, mainly composed by roots. Food stores contents analysis also provided information on the plants of interest for voles (*Airoldi 1976; Potapov et al. 2004)*. Other studies focused on indirect observations, such as resource depletion, to assess vole foraging behaviour in semi-captivity experimentation on prairie voles *(Cole and Batzli 1979*; *Kopp 1988*; *Saucy et al. 1999*, *Giban and Spitz 1967*) and common voles (*Spitz 1968*). All methods have shown their own advantages and limitations in diet interpretation.

To investigate the trophic niche of water voles, we also used several complementary methods based on DNA content analysis, determination of the species contained in food stores and behavioural observations. Due to the lack of methods to investigate the food preferences of subterranean species, we developed an original device with a camera trap to record the feeding behaviour of wild animals (*Lisse and Pinot 2024*) for which the first ecological results are presented in this study.

## Material and methods

### Fossorial water vole biology in the studied area

The water vole is known for its diversity in weight and body length, as well as its ecotypes (*Strachan et al. 2011*), ranging from a strictly aquatic form to a strictly fossorial form, with an intermediate form in between. Our study focused exclusively on the fossorial form of the water vole, *Arvicola amphibius*, in the Massif Central mountains in France. In this area, water voles inhabit exclusively natural grasslands used for farming production. The fossorial form of the water vole is characterized by its subterranean lifestyle, predominantly living in burrows and galleries, rarely surfacing except during sexual dispersal (*Airoldi 1978*; *Delattre et al. 2006*; *Saucy and Beat 1997*).

In the study area, the body weight of adult (i.e., reproductive) animal ranged from 71.2 and 137.6 grams (quantiles 0.05-0.95, N=2102 individuals; our data’s) and body length from 140 to 175 millimetres from the nose to the base of the tail. The populations exhibit strong amplitude cycles *(Pascal and Boujard 1987; Saucy 1988*) with periodicity ranging between 6 and 8 years (*Fichet-Calvet et al. 2000*).

### Study area and agricultural plots choice

The study was conducted in the natural habitat of the subterranean form of water voles, specifically in natural meadows managed by farmers. We selected numerous field plots within the study area (figure 1) using a stratified sampling. The plots were selected to reflect a gradient of elevation and a gradient of productivity. Productivity was estimated for each field plot within the study area by combining the administrative register of parcels and plant productivity from satellite image (HR-VPP products from Copernicus Land Monitoring Service by the European Environmental Agency and VITO, average value for years 2020, 2021 and 2022).

At the local scale (i.e., across several equivalent field plots), selection was based on technical considerations: road access, farmer’s knowledge and agreement, size of field plot (i.e., field plots smaller than 1.5ha were avoided), and the surrounding habitat of the plot (i.e., open field). In practice, potential fields plots were initially selected through GIS analysis (yielding dozens of equivalents for each level of elevation and productivity), then field-checked to ensure they met the criteria, and then asked for farmer agreement. The field plots selected (for each combination of elevation/productivity) were those for which farmers first provided their agreement, aligning with the random process inherent to stratified sampling.

The majority of the plots were located within outbreak areas. A minor part of the plots was in areas where water voles inhabit without exhibiting outbreaks.

### Methodological approach

The spatial and temporal structure of vegetation is crucial to understand herbivorous-plant interactions (*Soininen and Neby 2023*). Thus, we monitored the **available niche** (as defined by *Shipley et al. 2009*) accounting for seasonality and plant morphology (i.e. green parts VS roots).

To assess water vole’s diet, we described three levels;

First, we defined the **fundamental trophic niche** of the fossorial form of water vole. This was achieved using all available data on plant consumption: bibliography, single food tests in Campascopes (*Lisse and Pinot 2024*), roots found in food stores, DNA barcoding of vole stomach contents. We also added information on species eaten in market garden and orchards when we were able to catch voles on plant damage. This part of the study gave results on the diversity of potential diet at the species scale.

Second, we assessed the **realised trophic niche** in the meadows of our study area by considering data collected only from natural behaviours (food stores and DNA barcoding in stomach contents). This defined effective diet in meadows of our study area.

Third, we investigated the variability in **vole dietary preferences** by comparing the grassland composition through botanical, roots and DNA data (available niche) with the plants eaten/collected by voles (realised trophic niche). In addition, the variation in preferences was studied through triple food-choice tests conducted in Campascopes throughout the year on several plots.

Data use and concepts are summarized in figure 2.

**Figure 2:**
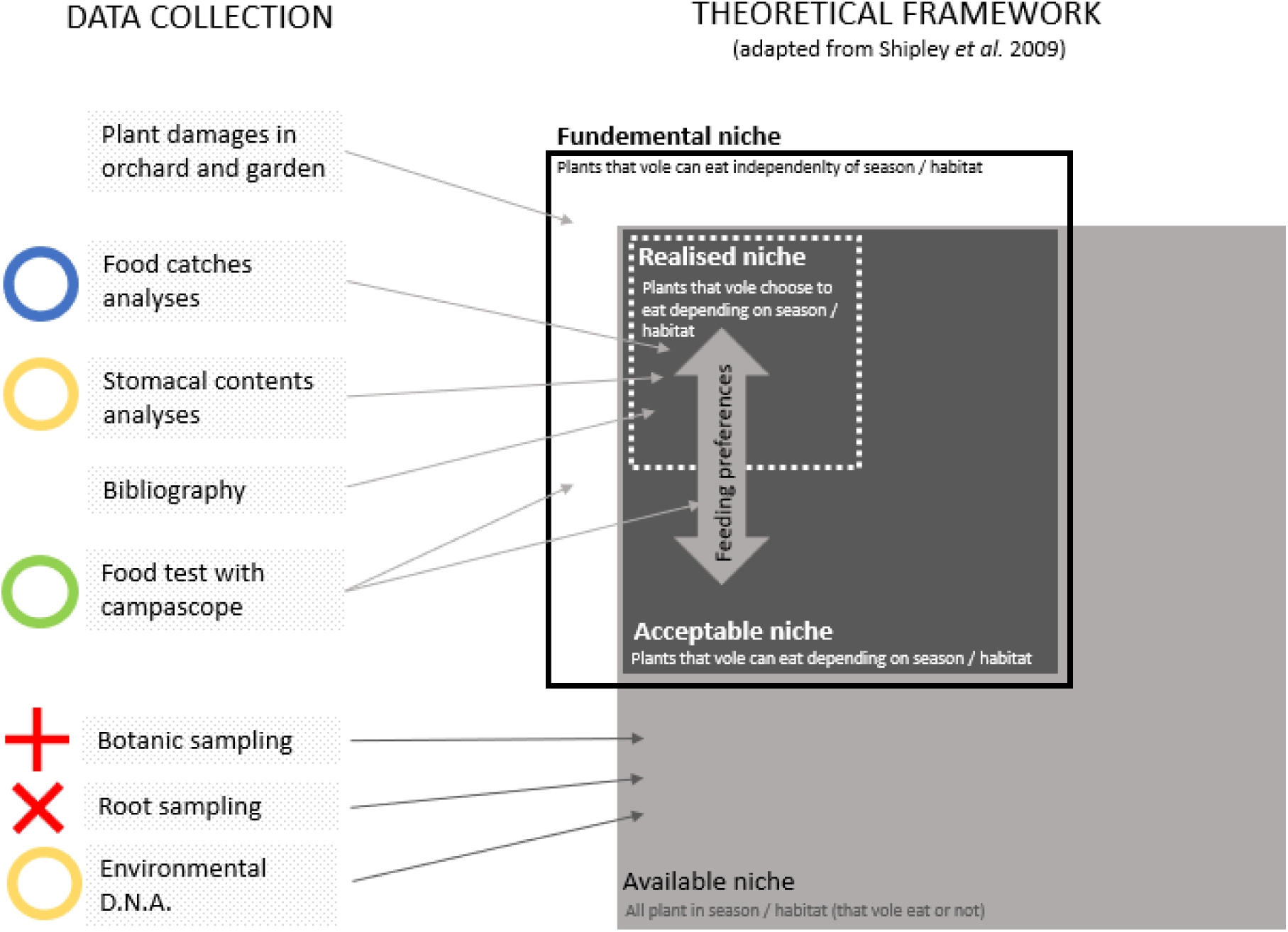
Overview of data collection and use. To investigate interaction between plants and fossorial form of water vole, different kind of data were collected (left) in natural habitat (cf. figure 1) and in bibliography. The link between data collected and the theoretical framework is highlighted by arrows. The theoretical framework is adapted from Shipley et al. (2009). Notice that there are no axes on ours.

### Data collection

#### Botanical data

To assess the available niche, plant species were inventoried in natural grasslands. Twenty-two field plots were sampled before vole outbreaks to avoid potential disturbances to the flora caused by vole overabundance (e.g. Nicod *et al.* 2020). The field plots were selected to represent a full gradient of productivity (figure 1), which is the main factor of vegetation variation in this area with the same climate. Three main types of meadows (Galliot *et al.* 2020) were sampled: MF25; Mountain hay meadow with *Umbelliferae* on healthy to fresh and fertile soil, MF26; Mountain hay meadow with *Rumex crispus* on healthy to fresh and very fertile soil, MP25; Mountain grazed meadow with *Lolium perenne* and *Taraxacum officinal*e on healthy to fresh and fertile to very fertile soil.

Samples were collected in spring 2019 or in spring 2021. For each plot, 9 random places of 0.33m² were described along a 100-meter transect in the middle of the grassland plot. Braun-Blanquet (1964) cover indices were noticed for each plant. The average proportion of each plant species was calculated.

#### Appetence of wild voles for meadows species using the Campascope

We developed an original design to propose food to voles inside their burrows. This device, the Campascope (*Lisse and Pinot 2024*), consists of a box buried in a gallery and equipped with a camera to film voles’ behaviour. We proposed a fresh small piece (∼20g) of each plant species found in meadows (list issued from botanical data and representing the available niche) and observed if the voles consumed it or carried it away. We also tested plants that water voles may not find in meadows but are consumed by other vole species (spruce, silver fire, *Jacob and Tkadlec 2010*) or were formerly used as livestock fodder in the study area (common alder, European ash).

#### Roots in grasslands and roots in food stores

To refine the assessment of the available niche, we also specifically estimated root proportions in grasslands. In march 2018, twenty-nine plots were sampled for root biomass. All plots were natural meadows used for hay production and were sampled during vole population growth phase. All these meadows revealed a high vole density during following winter (maximum observed density from 173 to 1179 individuals/ha, mean = 525, sd = 267). In each plot, roots were sampled with 10 cores along two intersecting transects, with 3 meters between each sampling point. The soil samples containing roots were collected using an 8 cm diameter root auger to a depth of 12 cm. The samples were then sieved using different mesh to separate roots by diameter and to remove soil. Root species were identified by visual inspection (*Taraxacum officinale, Trifolium sp., Ranunculus sp., Dactylis glomerata*, bulbs, and thin roots or other roots that could not be characterised to species/genus) and weighted.

From 2017 to 2021, we monitored *Arvicola amphibius* populations in these fields plot several times each year (with at least one trapping in spring and one in fall). During excavation for trap placement, we discovered 15 food stores across 8 different plots. For each food store, the contents were collected, cleaned, and the plant species identified. We also recorded the dates of findings to analyse the seasonality of food store discovery. For 6 stores of roots, each piece was weighed, measured and dried. To achieve this part, we supplemented our field observations with information from the literature (Airoldi 1976, Kopp 1998).

#### DNA material in grassland and in vole stomach

DNA metabarcoding analysis of vole stomach contents was compared to soil analyses. This comparison was conducted by sampling in summer on twenty-five plots of natural meadows with high vole densities. In each plot, ten adult voles were trapped using Topcat® lethal traps and dissected to extract their stomach contents. Concurrently, ten soil samples were collected with 8 cm diameter augers to a depth of 5 to 8 cm. Soil samples were made to assess vegetation diversity through eDNA, especially for plants that were not/few visible during botanical sampling (July) but could represent substantial resources due to their big roots). Soil samples and stomach contents from the same plots were pooled into single sample.

Before analysis, samples were prepared by scraping stomach contents and removing soil particles from plants, dried in accordance to Argaly laboratory protocols (see dedicated protocol in Apendix I), conditioned in 20 -gram sterile tea-bags with a calcium chloride additive and sent to the laboratory (Argaly, Society of DNA Analysis from environmental samples, located at Saint-Hélène-du-Lac -73800 FRANCE). All field and laboratory work were conducted under drastic hygienic condition (individual protection equipment and fire-cleaning of utensils) to avoid DNA contamination. DNA extraction and metabarcoding were made at Argaly laboratory according to procedure detailed in Appendix 1. DNA identification and the associated number of sequences were obtained for each batch of stomachs and soils samples. We first cleaned data provided by Argaly by eliminating matches with species and genus that were not local plants. Plants families were pooled to match identification between soils samples and stomachs contents. To avoid interpretation bias when analysing DNA sequences quantitatively, caused by preferentially amplified sequences (Taberlet et al. 2012; Pompanon, 2012), we compared the number of occurrences observed in the soils and in the stomachs.

#### Trapping on voles’ damages

To expend our knowledge on the fundamental trophic niche, we completed the list of eaten plants with opportunistic data collected from damages observations reported in the field by professionals (farmers, scientists) on cultivated plants. For each data, we confirmed the origin of the damage through systematic trapping. Data were collected in Massif Central mountains gardens and orchards from 2017 to 2023. Not all identified plants were systematically represented in natural grasslands.

### Statistical comparison between plant availability (available niche) in meadows and vole samples

Vole preferences were assessed by comparing the **realised trophic niche** with the **available niche**. The summer *selection coefficient* was estimated by comparing plants occurrences in stomach samples with plant frequencies in meadows (botanic: Braun-Blanquet and barcoding: occurrence of sample). The winter *selection coefficient* was similarly estimated by comparing the proportion of roots in food stores with the proportion of roots in meadows. These analyses were conducted at the family level These indicators provided a general pattern and were used to rank plants in order of importance for the vole diet. Shannon diversity indices (*Shannon 1948*) were calculated for each type of field data collected (botanical samples, stomachal DNA, soil DNA, roots in food stores, roots in soil). These indexes were used to evaluate the broadness of the different niches.

#### Test of feeding preferences variability in Campascope

To measure the representativity of food preferences pattern, the variability of these preferences regarding environmental factors of voles’ natural habitat was tested. Behavioural tests were conducted on wild voles in natural grasslands of both the study and control areas (Figure 1). These tests of feeding choices were carried out using the Campascope (*Lisse and Pinot 2024*) between three plants of interest. The plants were chosen from each major family group available in the field, based on species previously mentioned in the literature or identified as significant during the realised trophic niche analysis. Specifically, preference tests were conducted on a triptych of dandelion (*Taraxacum officinale*, noted as the preferred species), white clover (*Trifolium repens*, present in all grassland and also selected) and cocksfoot (*Dactylus glomerata*, present in all grasslands and apparently the most selected gramineous). We tested variability of food preferences between these 3 plants under different environmental conditions. For this, six representative grassland plots, each with variable plant composition and differing densities of the 3 species tested, were chosen. Four seasonal sessions were realised, in March (late winter), May (spring), June/July (summer) and September/October (fall). Because of snow cover and frozen soils, devices could not be set up during full winter. The densities of voles and dandelions were determined using remote sensing by analysing the amount of bare soil and the number of dandelion flower heads in drone-captured photos (see Buronfosse *et al.* in prep for full method description). Mean field plot productivity was estimated from vegetation index of sentinel data (total production of season one, resolution of 10m averaged at field plot scale, downloaded from Wekeo). A total of 128 Campascopes were set up, with at least 25 per season and 5 per plot. Fresh plants were collected directly from the field plot or its border. The three species *Trifolium repens*, *Taraxacum officinale* and *Dactylis glomerata* were each calibrated to be approximatively the same size and weight (∼15g) and randomly positioned in the box. When plants were all consumed, new fresh plants were placed in the Campascope. After the first choice, plants were left for a maximum of 1h30 (i.e., the standard length of a vole’s activity period during the day, *Airoldi 1979*), before resetting the choice with new fresh plants. The age class of the vole (juvenile, subadult or adult) was estimated from each video based on its size *(Saucy 1988*). For each plant species, the rate of removal or consumption was recorded, as well as the order in which removal or consumption occurred.

Generalised linear model (Poisson) was employed to estimate the variation in average choice rank for each of 3 plants tested. This model enabled us to determine whether significant differences existed between the 3 plants. The influence of external variables on the ranking was tested with sub-models for each plant (Table 2). The following variables were tested : season (factorial variable; spring, n=25; summer, n = 56, fall, n = 27, winter, n = 32), elevation (numerical value, [530, 1100]), average meadow productivity (numerical index data obtained from sentinel, [1000, 2475]), apparent vole density (factorial variable, class of density of earth mounds : low, middle, high), dandelion density (average number of flower per 0.58 hectare recorded in field plot on May 15^th^ by remote sensing, numeric variable, [0, 140 155]) and individual age class (based on visual estimation of size from videos recordings, 3 classes : juveniles, subadults and adults). Each variable was tested independently and results compared with the null model using AIC comparison. Models with a delta AIC < 2 were assumed equivalent models and the best model was selected as the one with the fewest parameters considered (*Burnham and Anderson 2002*).

All applicable institutional and national guidelines for the care and use of animals were followed. This study was conducted in accordance with the guidelines declared in the Declaration of Helsinki, and received approval from the competent authorities (no. 37713, project authorization no. 2238) and from the ethics committee of the veterinary school of Lyon.

## Results

### Fundamental trophic niche

Water voles were found to consume at least 79 different plant species, belonging to 30 different plant families. These species are listed in table 1, specifying the type of data recorded and their presence in the meadows within our study area. The water vole did not exhibit neophobic behaviour, as it consumed leaves of forest trees. It consumed also plants with toxic components (e.g. *Narcissus*, *Rumex*, *Ranunculus*) as well as those with urticant properties (*Urtica*). Voles were able to eat a large variety of herbaceous plants, but also roots of creepers (2 species) and trees (6 species). Voles also ate plants formerly used for feed livestock (common alder, European ash) but ignored resinous trees (spruce, silver fire). Voles were able to eat all the available plants in grasslands.

**Table 1:** Plants species eaten by fossorial form of water vole, *Arvicola amphibius.* Habitat represent the main habitat of the species (G = grasslands, M = market-gardening, C = cultures, O = orchards). Braun-Blanquet represent the abundance-dominance index observed in our study area. The next 5 columns are the results of feeding observation (a cross means that the species was used/eaten by vole in that observational condition).

| Species | Habitat | Braun-Blanquet | Damages | Food stores | Stomach | Campanoscope | Koop / Airoldi papers |
| --- | --- | --- | --- | --- | --- | --- | --- |
| *Achillea millefolium* | G | 1 |  |  |  | x |  |
| *Agropyrum repens* | G | / | x | x |  |  |  |
| *Allium ampeloprasum* | M | / | x |  |  |  |  |
| *Alnus glutinosa* |  | / |  |  |  | x |  |
| *Alopecurus pratensis* | G | 1 |  |  |  | x |  |
| *Anethum graveolens* | M | / | x |  |  |  |  |
| *Anthriscus sylvestris* | G | 0 |  |  |  | x |  |
| *Avena sterilis* | G | / |  |  | x |  |  |
| *Bellis perennis* | G | / |  |  |  | x |  |
| *Beta vulgaris subsp. Vulgaris* | M | / | x |  |  |  | x |
| *Bromus hordeaceus* | G | 1 |  |  |  | x |  |
| *Capsella Bursa-pastoris* | G | 0 |  |  | x | x | x |
| *Castanea sativa* | O | / | x |  |  |  |  |
| *Cerastium arvense* | G | 1 |  |  |  | x |  |
| *Cichorium intybus* | M | / | x | x | x |  |  |
| *Cirsium arvense* | G | 0 | x |  |  | x |  |
| *Convolvulus arvensis* | G | / |  |  |  |  | x |
| *Crocus albiflorus* | G | / |  |  |  |  | x |
| *Cucurbita pepo* | M | / | x |  |  |  |  |
| *Cydonia oblonga* | O | / | x |  |  |  |  |
| *Cynosurus cristatus* | G | 0 |  |  |  | x |  |
| *Dactylis glomerata* | G | 6 |  | x |  | x | x |
| *Daucus carota* | G | 0 |  |  | x | x | x |
| *Daucus carota subsp. Sativus* | M | / | x |  | x | x | x |
| *Euphorbia cyparissias* | G | / |  |  |  | x |  |
| *Festuca arundinacea* | G | 2 |  |  |  | x |  |
| *Festuca rubra* | G | 1 |  |  | x | x |  |
| *Fraxinus excelsior* | O | / |  |  |  |  |  |
| *Galium verum* | G | 0 |  |  |  | x |  |
| *Heracleum sphondylium* | G | 2 |  |  |  | x | x |
| *Halcus lanatus* | G | 5 |  |  |  | x |  |
| *Humulus lupulus* | C | / |  |  |  |  | x |
| *Knautia arvensis* | G | 1 |  |  |  | x |  |
| *Lactuca sativa* | M | / | x |  |  |  |  |
| *Lathyrus pratensis* | G | 0 |  |  |  | x |  |
| *Leontodon hispidus* | G | / |  | x |  |  |  |
| *Leucanthemum vulgare* | G | 2 |  |  |  | x |  |
| *Lolium perenne* | G | 9 |  |  |  | x | x |
| *Lotus corniculata* | G | 0 |  |  |  | x |  |
| *Malus domestica* | O | / | x |  |  |  |  |
| *Medicago Sativa* | C | 0 |  |  |  | x | x |
| *Myosotis arvensis* | G | 0 |  |  |  | x |  |
| *Narcissus pseudonarcissus* | G | 0 | x |  |  |  |  |
| *Petroselinum crispum* | M | / | x |  |  |  |  |
| *Phleum pratense* | G | 0 |  |  |  | x | x |
| *Plantago lanceolata* | G | 3 |  |  | x | x |  |
| *Plantago major* | G | 0 |  |  | x | x |  |
| *Poa pratensis* | G | 8 |  |  |  | x |  |
| *Poa trivialis* | G | 8 |  |  | x | x |  |
| *Potentilla erecta* | G | 0 |  | x |  | x |  |
| *Pyrus communis* | O | / | x |  |  |  |  |
| *Ranunculus acris* | G | 4 |  |  |  | x |  |
| *Ranunculus bulbosus* | G | 0 | x | x |  | x |  |
| *Ranunculus repens* | G | 0 |  | x |  | x |  |
| *Rhodiola rosea* | M | / | x |  |  |  |  |
| *Rumex acetosa* | G | 2 |  | x |  | x | x |
| *Rumex crispus* | G | 1 |  |  |  | x |  |
| *Sanguisorba minor* | G | 0 |  |  |  |  | x |
| *Scorzonera autumnalis* | G | 0 |  |  |  | x |  |
| *Silene vulgaris* | G | 0 |  |  | x |  |  |
| *Solanum tuberosum* | M | / | x |  |  |  | x |
| *Spinacia oleracea* | M | / | x |  |  |  |  |
| *Stachys officinalis* | G | 1 |  |  |  | x |  |
| *Stellaria media* | G | 2 |  |  |  | x |  |
| *Taraxacum officinale* | G | # | x | x | x | x | x |
| *Tragopogon pratensis* | G | 0 |  |  | x |  |  |
| *Trifolium dubium* | G | 1 |  |  |  | x |  |
| *Trifolium pratense* | G | 1 |  |  | x | x | x |
| *Trifolium repens* | G | # | x | x | x | x | x |
| *Trisetum flavescens* | G | 5 |  |  | x | x |  |
| *Tulipa sp* | M | / | x |  |  |  |  |
| *Urtica dioica* | M | / | x |  |  |  |  |
| *Veronica arvensis* | G | 0 |  |  |  | x |  |
| *Vicia faba* | M | 2 | x |  |  | x |  |
| *Vicia sativa* | G | / |  |  |  | x |  |
| *Vicia sepium* | G | / |  |  |  | x |  |
| *Viola tricolor* | G | 0 |  |  |  | x |  |
| *Vitis vinifera* | O | / | x |  |  |  |  |
| *Zea mays* | G | / |  |  |  | x |  |

**Table 2:** AIC selection tables for set of models explaining rank choices of plants proposed in behavioural test. (one model per variable tested/per specie). Selected model is in bold.

| Model set | Null model | Season | Elevation | Productivity | Vole density | Dandelion density | Vole age |
| --- | --- | --- | --- | --- | --- | --- | --- |
| Dandelion rank | **319.38** | 325.35 | 321.09 | 321.36 | 322.99 | 321.93 | 322.22 |
| Clover rank | **317.54** | 322.44 | 318.71 | 319.48 | 320.44 | 321.47 | 320.84 |
| Cock's-foot rank | **315.34** | 320.01 | 316.58 | 316.05 | 317.81 | 318.06 | 319.30 |

Fossorial form of water vole showed a very large fundamental trophic niche with no specific specialisation for the flora of meadows.

### Realised trophic niche

#### DNA in stomachs

DNA barcoding analyses of stomach contents identified 11 families of plants representing at least 20 different species. The Shannon index of diversity for the all dataset was 4.15, indicating a large and equilibrate diet *in natura*, at least in summer. During this period, 71% of stomachs contained *Poacae*, 71% *Asteraceae (*mainly *Taraxacum officinale)*, 39% *Polygonaceae* (mainly *Rumex sp*), 32% *Fabaceae*, 23% *Plantaginaceae*, 16% *Brassicaceae* and 13% *Caryophylaceae (Figure 3B),* highlighting the dominant role of *Poacae* and dandelion in the diet, but also the significant contribution of other plant families. The average Shannon index per vole was about 0.84 +/- 0.71 (mean +/- SD). In comparison with the Shannon index of all dataset, it suggests that vole diet is variable between individuals.

**Figure 3:**
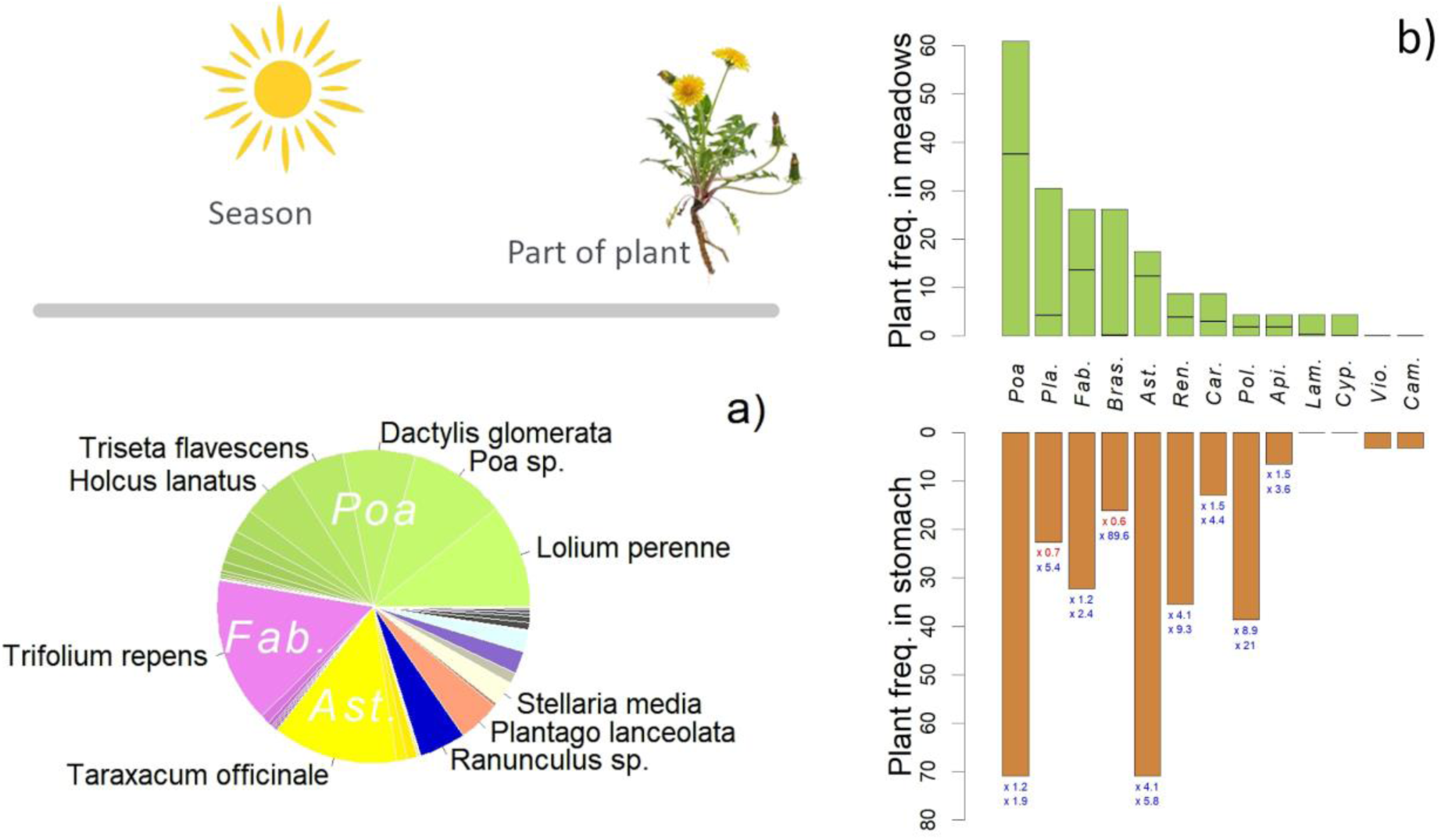
a) Grasslands composition in late spring from botanical records. b) Grasslands composition in summer (after mowing) from soil DNA analyses (green) and plant species distribution in vole stomach at the same time (brown). Horizontal bares in the green part of the graph represent occurrence of the family observed in the botanical sample (Braun-Blanquet). Below the brown bar plot, 2 coefficients are printed. The upper one is the comparison between DNA occurrence in soil (green bares) and in stomach (brown bares). The lower one is the comparison between Braun-Blanquet occurrence (horizontal bares) and DNA occurrence in stomach (brown bares). *Abbreviations; Pla.: plantaginaceae, Fab.: fabaceae, Brass.: brassicaceae, Ast.: asteraceae, Ren.: renonculaceae, Car.: caryophyllaceae, Pol.: polygonaceae, Api.: apiaceae, Lam.: lamiaceae, Cyp.: cyperaceae, Vio.: violaceae, Cam.: Campanulaceae*

#### Food stores

Over the course of 184 capture sessions and almost 1,000 colonies sampled, 15 different food stores were discovered, exclusively in autumn and winter, during the non-growing period.

Food stores were composed exclusively with roots cut into pieces and of low diameters. Among the 1596 fragments measured from 6 different food stores, a bimodal distribution was observed with short fragments (4,4 +/- 1.5 cm; 8.4 mm of average diameter) and long fragments (7.5 +/- 2.6 cm ; 8.2 mm of average diameter). The pieces of roots were well-preserved, as voles isolated them from each other by surrounding each fragment with soil. The weight of the food stores (excluding soil) was comprised between 37 and 1380 grams, with a mean of 545 grams.

The food stores contained plant material from 4 families representing 5 species (*Figure 4B*). The Shannon index of diversity was 0,366, indicating a low level of diversity. This suggests a highly constrained realised trophic niche for food stores. Indeed, 94.5% of the food stores were composed of dandelions roots, revealing a strong dependence on dandelions in winter in our study area.

**Figure 4:**
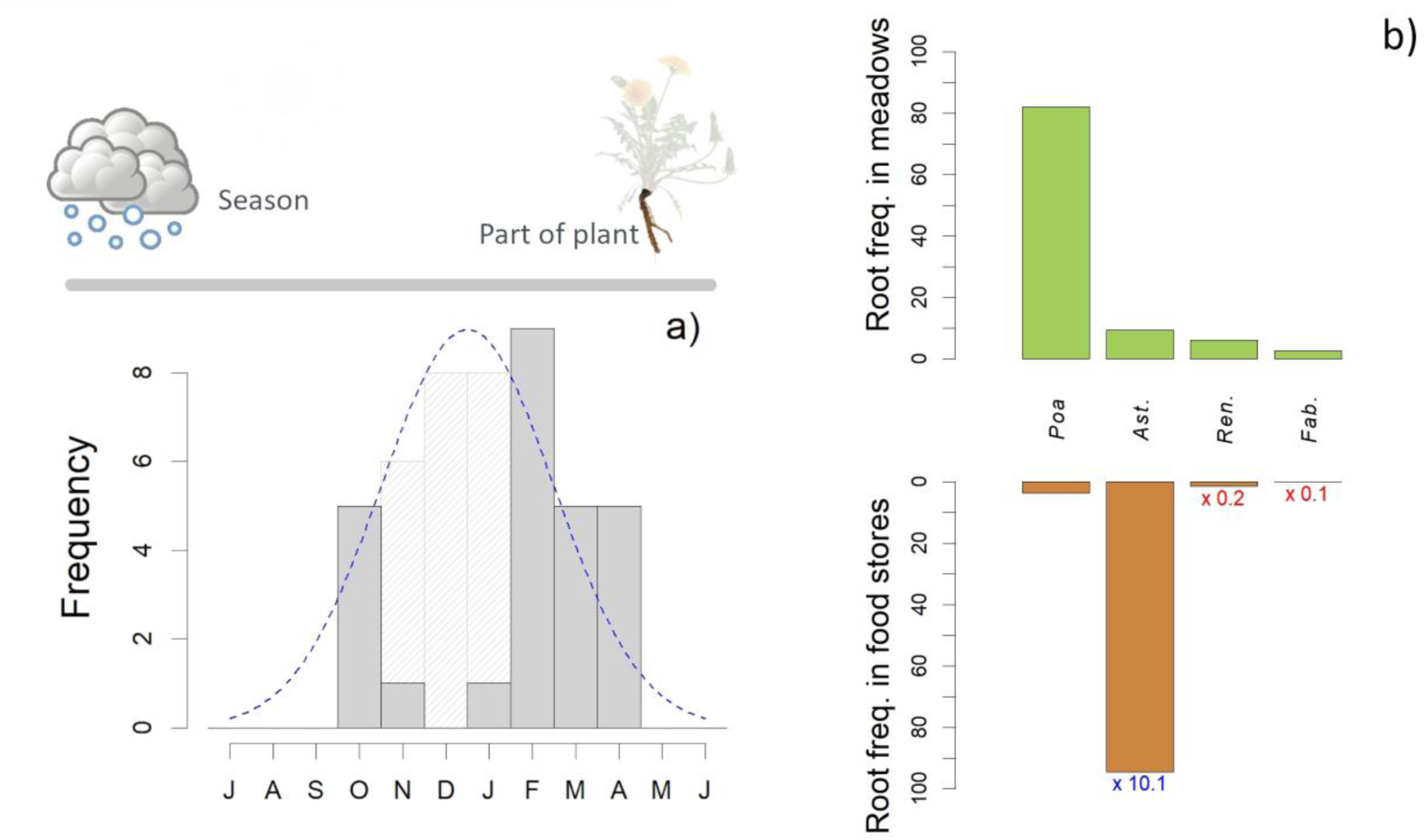
a) Seasonality of vole food stores discovery. Grey bares represent food stores observed in bibliography and recorded on field, light bares represent the expected food stores undiscovered due to lack of field work (due to snow and frozen soil). The blue line represents the expected seasonal kinetic of food stores. **b) Roots species available in grasslands during winter (green) and roots species found in voles food stores.** Below the brown bar plot, the comparison between root occurrence in grasslands and food stores is printed. *Abbreviations; Fab.: fabaceae, Ast.: asteraceae, Ren.: renonculaceae*.

### Food preferences

#### Plant availability: grasslands composition and diversity

At the community scale, we observed that vole inhabits highly diverse meadows. A total of 66 plant species were identified before mowing. Figure 3A shows the average plant composition of the 22 sampled grasslands located in outbreak areas (measured before outbreak to avoid vole disturbance). The average cover percentage was calculated for each specie (mean of percentage cover across 22 grasslands) and permitted to better describe the habitat of vole: grasslands were dominated by grasses, which accounted for approximately 40% of the cover and included 19 species (Appendix 2). The most prevalent Poaceae was *Lolium perenne* with 8.5% of cover, followed by *Dactylis glomerata* (5.9%) and *Triseta flavescens* (4.61%). Fabaceae were also strongly represented, accounted for 13.6%. *Trifolium repens* was the most abundant species, followed closely by *Taraxacum officinale* (10.3%). Other notable species included *Plantago lanceloata* or several *Ranunculus*. The Shannon index for plant diversity of grasslands was 4.13.

Soil samples for DNA analysis were collected after mowing and revealed the presence of 11 families of plants representing at least 22 different species. The Shannon index of diversity for the soil samples was 3.81, which is the same order of magnitude to the Shannon index obtained from botanical samplings. The dominant family identified was *Poaceae* present in 61% of the samples. Other prominent families included *Plantaginaceae*, *Fabaceae* and *Brassicaceae* (found in 30%, 26% and 26% of the samples, respectively). The *Asteraceae* family was present in 17% of the samples. Those results can be compared to the botanical results in fig 3B, though correspondence may be challenging due to potential inaccuracies in the recognition of DNA sequences. The relative contribution of plant families to the overall plant community was globally consistent between methods except for two families; *Plantaginaceae* and *Brassicaceae* which were over-represented in DNA soil samples compared to botanical samples.

In the soil, root biomass was dominated by root hairs from graminoids or other plants that could not be visually identified in winter (Figure 4A). They accounted for 61.1% of the total dry weight. The remaining biomass was composed of roots from *Dactylis glomerata* (20.8%), *Taraxacum officinale* (9.4%), *Ranunculus sp.* (6.1%) and *Trifolium repens* roots (2.6%). The Shannon diversity index for root biomass was 1.61. This low value should be interpreted with caution because of the great difficulty in identifying small root species.

#### Preferences observed in DNA analyses

The Shannon diversity index for vole stomach samples was about 4.15, higher than that observed for soil DNA samples but similar to the index obtained on botanical samples. However, the diversity in vole stomach contents was mainly due to diversity between individuals rather than diversity within the same bolus. Indeed, the average Shannon index of diversity for an individual was about 0.84. Despite this, it is unlikely that individuals were specialised as the average species richness per stomach was 5.22 (SD : 3.52). The most selected family seemed to be *Polygonaceae* (mainly *Rumex acetosa)* with an 8-fold increase compared to soil DNA and a 21-fold increase compared to botanical samples, then *Ranunculaceae* (4.1 to 9.3-fold increase, probably *Renunculus bulbosus*) and *Asteracaea* (4.1 to 5.8-fold increase, *Taraxacum officinale*). Some other families also seemed to be common in stomachs, despite being relatively rare in meadows; the *Caryophylaceae* (1.5 to 4.4-fold, mainly *Silene vulgaris*), the *Apiaceae* (1.5 to 3.6-fold, mainly *Daucus carota*). Voles also appeared to select a *Brassicaeae* (*Capsella bursa-pastoris*) that was not detected in botanical samples. While the voles’ stomach contents consisted mainly of *Poaceae*, it was unclear whether selection had occurred, as the family was highly dominant in grasslands. *Fabaceae (*mainly *Trifolium sp)* and *Plantaginaceae* (*Plantago* sp) were probably in the same case. None of the dominant plants were avoided. Because comparisons between botanical and environmental DNA samples showed that *Plantaginaceae* and *Brassicaceae* were overrepresented in DNA soil samples compared to botanical samples, quantitative results from DNA stomach samples should then be taken with caution. With this caution, a pattern of selection in summer is found for *Rumex*, *Ranunculus* and *Taraxacum*. The common feature of the apparently selected species is a taproot.

#### Preferences observed in food stores

Food stores were composed of 94.5 % of dandelion roots (*Taraxacum officinale*), even though this species accounted for only 9.4 % of the available root biomass in grassland samples (Fig. 4B). The remaining 5.5% comprised little bulbs, which were absent from the soil samples, indicating that voles are likely to find and select them, and of *Trifolium sp* and *Ranunculus sp*, which were also found in the soil. When comparing the proportions of plant families between food stores and soil samples (fig 3B), all families were markedly underrepresented, except for *Astearace* (10-fold increase), indicatinga strong selection of *Taraxacum officinale* by water voles.

#### Variability of food preferences

In the field, we recorded vole food choices on video throughout several seasons on an entire year. After visioning records, 140 sequences of vole choice, involving at least 70 animals, divided into 4 batches of 20 to 42 videos (1 batch per season) were available for statistical treatment (Table 2). Average choice ranks for *Taraxacum officinale, Trifolium repens* and *Dactylis glomerata* were 1,42 +/-0.106, 1.77 +/-0.122, 2.6 +/-0.159, respectively (+/- are standard errors from the Poisson model, Figure 5). Voles showed a high preference for *Taraxacum officinale* which was taken in 95% of cases and chosen first in 60% of cases. *Trifolium repens* was taken in 85% of cases and chosen first in 32% of cases, while *Dactylis glomerata* was taken in 75% of cases and chosen first only in 8 % of cases.

**Figure 5:**
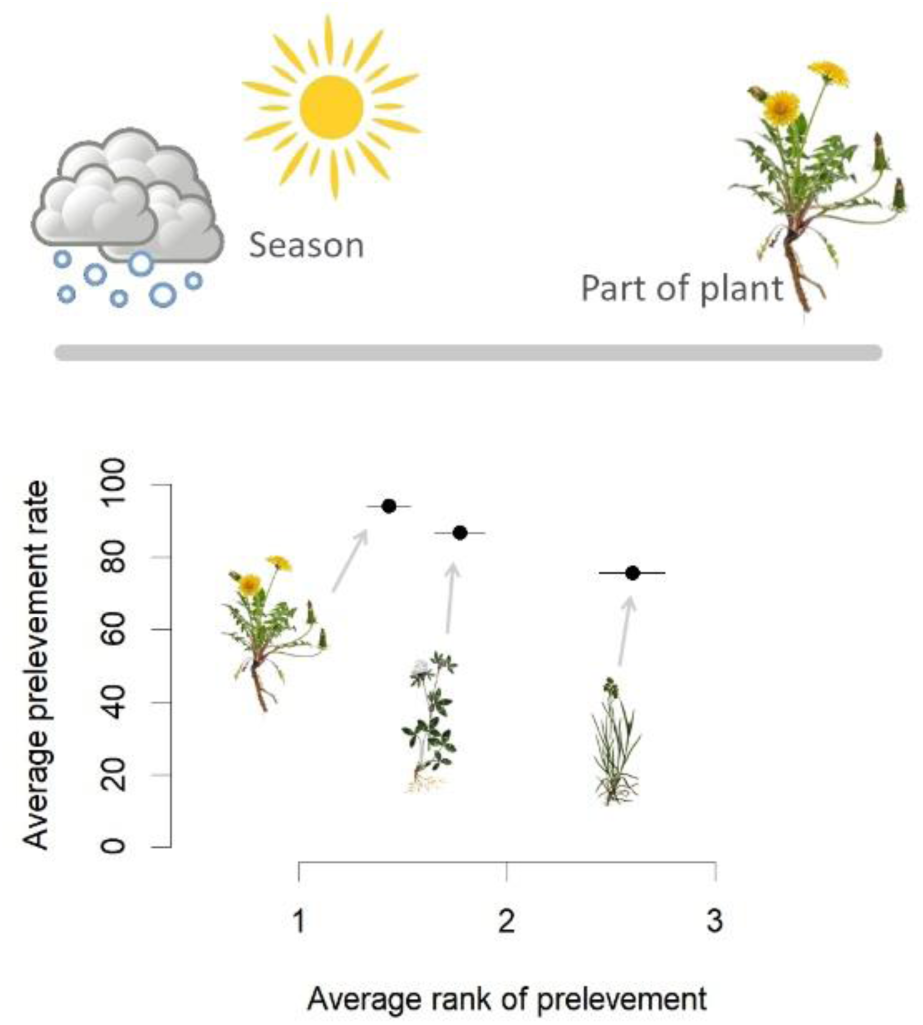
Average percentage of plants that voles eat or bring as a function of their average choice ranks. The three same plants (*Taraxacum officinale*, *Trifolim repens*, *Dactylis glomerata*) were systematically proposed at the same time during behavioral tests. Behavioral tests were spread along environmental gradients and over the course of seasons.

None of the tested factors (seasons, elevation, average grassland productivity, apparent vole density, dandelion density, individual age class) significantly influenced the explained variable (i.e., choice rank of each tested plant) according to the AIC selection model procedure (table 2). The best-fitting model in each case was the null model, which only estimated the average choice rank for each plant (as shown above or in Figure 5 for estimated coefficients and standard errors). This result highlights the very low variability in voles’ food preferences.

## Discussion

### Trophic niche of water vole: consistency of general pattern

#### Investigating fundamental trophic niche

While the literature listed several plants consumed by water voles, the use of the Campascope device has expanded our knowledge of the plants species likely to be consumed by them. Conducting feeding choice tests within the natural habitat of water voles appeared to be a good way to collect information about wild animals, as captivity is known to introduce biases in behavioural responses, especially for specialist species *(Shipley et al. 2009).* Although the list of accepted species is not exhaustive, the 79 different plants tested in this study demonstrated that water voles have a broad fundamental niche, including plants not found in grassland habitats.

#### Investigating diet during growing season

DNA analysis of stomach contents was the main tool used to investigate diet during the growing season, providing new data on the diet of water voles in their natural habitat. Over the past decade, advancements in DNA extraction techniques for environmental studies has enabled the description of the diets of many cryptic wild species in the field, especially rodents (*Soininen et al. 2009*, *Neby et al. 2024*). Faecal samples are also a common way to investigate rodents’ diet (e.g. *Neby et al. 2024*) because it is non-invasive. However, faecal samples proved complicated to carry out for water voles without strong animal disturbance and intensive field work effort. Indeed, water voles are strictly subterranean and also have subterranean latrines, which complicates access to these biological materials. These also deteriorate very quickly when exposed to the dampness of the burrows. Stomach sampling was then the best option to work with less degraded materials.

In this study, DNA analysis provided valuable insights into the fundamental trophic niche, consistent with the results of feeding tests on voles in the Campascope device. Water voles actually feed on a wide range of vegetation.

However, the DNA metabarcoding of stomach contents in this study exhibited several biases that could alter the quantitative interpretation of the data, especially in estimating the realised trophic niche. Indeed, DNA sequencing lacked precision for some taxa and the DNA database used for DNA comparison might have influenced species determination (*Pompanon et al. 2012*). To compensate for this bias, the precision of the data had to be degraded to the taxonomic level of family. Although this allowed a very consistent pattern to be drawn, it was unsatisfactory for families such as *Fabacea*, which are thought to be important for voles and encompass several known species. Additionally, stomach content analyses may be biased by over-representation of the last meal, particularly in species such as voles, where food intake is limited by stomach size (*Zynel and Wunder 2002*). To compensate for some of the last meal effect, the number of occurrences in overall stomachs was used rather than the total number of sequences.

#### Investigating diet during the winter season

Assessing the winter diet and available winter niche is challenging but crucial for understanding rodent-plant interactions (*Soininen and Neby 2023*). Analysis of food stores has provided valuable data into vole behaviour, although such data remain opportunistic and difficult to obtain. The field effort required to discover food stores could not be precisely quantified; however, only 15 food stores were discovered in 184 trapping sessions. Although trapping was conducted throughout the year, we only found food stores during the non-growing season (from middle autumn to early spring). We also observed a peak in October and another in February to April. However, it is possible that the number of food stores in November, December and January was significantly underestimated due to reduced trapping during these months associated to frozen ground and snow cover. Nevertheless, the seasonal pattern seems to be consistent despite the relatively small sample size.

In our study area, dandelion was almost the only species stored by voles, which is consistent with their food preferences observed during the growing season. However, we cannot assert that voles did not consume other plants that they did not store. Airoldi (1978) reported more diverse food stores, indicating that the food storage behaviour of the water vole in our outbreak area is specific to dandelion, but it might be more diverse in other areas.

#### Investigating variability of food preferences

To assess feeding preferences in the wild, the Campascope device provided accurate data, considering the extrinsic and intrinsic potential factors of variation. This method provided good quality behavioural data. However, this device does not allow many plants to be tested simultaneously and requires significant field investment. To optimise fieldwork, we relied on previous literature, DNA analysis of stomach contents and analysis of food store contents to select three plants (the three most selected plants of the three most important families). These plants were then test to examine seasonality in voles’ food preferences and cultural habits. Our results indicate minimal variability with dandelion consistently emerging as the preferred plant. This approach appeared necessary to complement other samplings methods, as the study investigate strong plant-rodent relationship. Indeed, preferences may be overlooked due to unperfected reflection of preferences in plant availability, especially during the winter season. On the contrary, seasonal shift of diet may not be observed due to voles’ choice of settlement site with sufficient resources for its overall life season. In addition to the contents analysis that guided plants to test in the cafeteria test, the direct choice tests were conducted across different seasons allowed to avoid these two situations.

#### The general pattern

This study identified a general pattern for the diet of water voles in outbreak area:

- Water voles have a very broad fundamental niche.
- Their feeding behaviour displays a strong seasonal pattern, with food stores being used during the winter season and probably only direct intake during the growing season.
- During the growing season, voles have a large realised trophic niche.
- In the winter season, the realised trophic niche seems to be very smaller than that observed during the growing season and voles appear to be constrained by their reliance on dandelions for food stores.
- Voles consistently prefer dandelions regardless of environmental or intrinsic variations.

The fossorial water vole is therefore probably a facultative specialist, as defined by Shipley and co-authors (2009). While its realised niche is broad during the growing season, it becomes narrower in winter (Figure 6). During this period, voles appear to forage on roots, a difficult resource requiring morphological adaptation for burrowing. However, the water vole is clearly not a specialist, as it shows a broad realised niche in summer and does not specialise in difficult chemical plants: it is able to eat *Ranunculus* or *Rumex*, which also have large roots, but does not show strong selection for them in winter. Furthermore, it is obvious that water voles are able to live in few productive areas (*Fichet-Calvet et al. 2010*), where dandelions are scarce.

**Figure 6:**
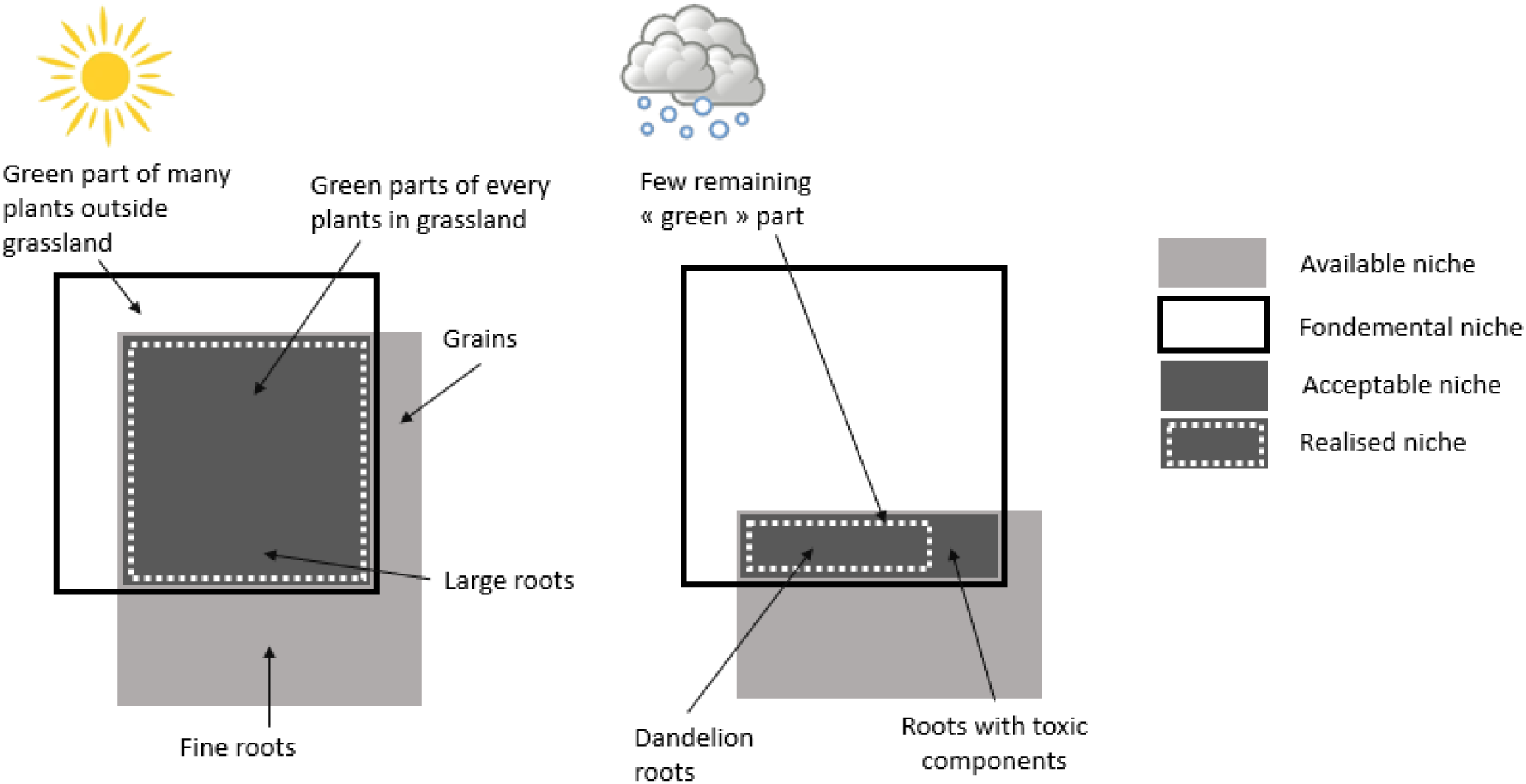
Conceptual schematisation of water vole niches through the seasons. The realised niche of water vole seems to be highly constrained during winter by the availability of resource. It results in a strong specialisation on dandelion on our study area during that season, but water voles showed a preference for dandelion all around year.

### Implications for population cycles and plants hypothesis

The water vole’s large fundamental trophic niche enables it to survive in various types of grasslands, ranging from low-to highly-productive meadows, from the Iberian Peninsula to Siberia. However, while water voles thrive in some meadows, they are rare in others.

There is ongoing debate regarding whether water voles exhibit population cycles. The main reason for this is the lack of long-term series, as the expected cycle for an animal of this size is 5 to 8 years (*Peterson et al. 1984*). However, Saucy (1994), based on data from a large-scale control programme (“tail return statistic”), showed that regular cycles in vole numbers, lasting between 5- and 7-years, occur at the municipal scale in most of the surveyed municipalities. Korpimäki *et al.* (2005) reported 8 to 10 years cycles in Western Finland. Those studies were conducted far from our study area. In our study area, vole outbreaks have been monitored by the government since the 1980s. Unfortunately, the observational process was not designed for scientific analyses (the records of vole densities are of low resolution and the grain of survey has changed 3 times in 40 years). However, these surveys have revealed regular cycles, synchronous over a very large scale (*Note and Poix 2006*), with periods of 6 to 8 years.

In our effort to understand cyclic dynamics, we focused on the water vole in the French mid-mountains as a candidate system to test the plant-hypothesis.

To evaluate whether a rodent cyclic system is driven by the plant-hypothesis, three main points have to be checked: i) Plants show strong seasonality in their growth period, ii) Rodents have strong food preferences or dependence on a small number of plant species, iii) After rodents have damaged their preferred food sources, a recovery period is needed to restore its quality or quantity.

Before delving into further investigations, the diet of this species had to be elucidated. The results of the present study on the water vole’s trophic niche in the French mid-mountains provided insights into the interest of this plant-hypothesis for further understanding this cyclic system.

### 1/ seasonality of trophic niche depending on plant growing season

As no water vole cycles have been documented in oceanic climates, seasonality likely plays a crucial role in the occurrence of water vole cycles. Investigating the trophic niche across seasons in a mid-mountain climate provided insights into the seasonality of vole-plant interactions. Under these seasonal conditions, the vole’s available niche differs significantly between the growing season (broad available niche) and winter (narrow available niche). As a generalist species, water voles seem to be well-adapted to all grasslands during the growing season but are less so in winter.

Indeed, we found that water voles have a broad fundamental trophic niche that allows them to feed on every plant species in the grassland. In spring, summer, and early autumn, grasslands produce a high quality and quantity of plants, providing voles with abundant food and resulting in a broad realised niche. As the grasslands in our study area are devoted to livestock farming, most farmers tend to maintain high productivity in their plots. During the growing season, voles are probably more dependent on grassland productivity than on specific plant composition.

In our study, water voles select plants with taproots during the growing season, such as *Taraxacum officinale*, *Rumex sp*, and *Ranunculus sp*. Their diet is also completed by *Poaceae*, which are widespread in the studied grasslands, including species like *Dactylis glomerata*, *Poa trivialis*, and *Tristemum flavescens*. Additionally, voles consume clovers. The preference of voles for protein-rich species, such as clovers, vetches, or alfalfa, has been assessed in previous studies on other vole species (*Bergeron and Jodoin 1987*, *Cole and Batzli 1978*, *Curtis et al. 2002*, *Marquis and Batzli 1989*, *Dejaco and Batzli 2013*). These preferences are consistent with previous research on the diet of *Arvicola amphibius* in Switzerland (*Saucy 1988*, *Saucy et al. 1999*). However, as clovers are abundant, they do not appear to be over-selected. Saucy *et al.* (1999) noted that clover may contain a cyanogenic component that makes it toxic and limits its intake.

Almost all grasslands are grazed by livestock in the fall, leaving only lawn-height vegetation for the winter. As a result, the amount of food available to herbivores decreases drastically in winter, making rodents highly dependent on food storage behaviour. Such storage behaviour is well-documented in several rodent species and has been selected in evolutionary processes to survive the winter season (e.g. *Soulairac and Lapetite 1946* review on rats; *Potapov et al. 2004*, on water voles). In our study, the storage behaviour of water voles was only observed in winter, suggesting their dependence on these food storages to survive the winter season (*Potapov et al. 2004*). In our study area, food storage of water voles seems to be mainly dependent on only one species, the dandelion. The low proportions of *Rumex sp.* and *Ranunculus sp.* in the food storages may be explained by the presence of toxic components, oxalic acid and protoanemonin respectively, as well as by their lower densities in our meadows. The voles’ realised trophic niche seems to be narrower in winter, as a result of the narrowest available niche.

### 2/Strong feeding preferences for dandelion

If dandelion is the only choice in winter, it should be noted that it is also the first choice in summer, even in meadows where there are no dandelions, and is preferred to clover and cocksfoot in all seasons. This suggests an innate behaviour. The interest of dandelion for rodents seems obvious because of its large, non-toxic taproot. Furthermore, the strong preference of voles for dandelion may provide a significant advantage in seasonal environments.

Previous studies have also found a feeding preference for dandelion in the montane vole (*Marquis and Batzli 1989*, *Dejaco and Batzli 2013*), but in third place after *Medicago sativa* and *Trifolium pratense*, and in the Orchogaster vole (*Curtis et al. 2002*). Dandelion has also been noticed as a strongly selected species by the water vole in Switzerland (*Kopp 1988*, *Airoldi 1976*).

### 3/ Preferred resource depletion and resource recovery delay

In line with the maximum number of dandelion apexes found in a food store (233) and the local density of dandelions on the same plot (27 individuals per square meter in uncolonized areas), the storage behaviour of voles probably may significantly impact the local density of dandelions: a food store may contain all the dandelions from an area of 8.5 square metres for a vole territory size of around 50 square meters.

In fact, in our study area, dandelions can be very abundant in grasslands, with more than 10 plants per square meter, representing a potential resource of 500 grams per square meter. However, observations of root fragments in food stores indicate that dandelions are unlikely to regrow after harvest. Given the unlikelihood of replenishment of the dandelion root bank within a single year, there is a risk of depletion of the resource.

Thus, given the current state of knowledge, the plant hypothesis for explaining water vole cycle dynamics in the mid-mountains should be seriously considered.

To further test this hypothesis, it will be useful to investigate i) whether voles affect the quantity or quality of dandelions, and if in turn, whether dandelion deficiencies affect the reproduction, survival or dispersion of voles ii) whether the dandelion population regrows or restores its intrinsic quality after being heavily foraged, and with which delay.

## Conflict of Interest

We declare no conflict of interest, that all authors of the manuscript have seen and approved the submitted version, that all authors have contributed substantially to the work, and that all co-authors have been included.

## Funding

The study was funded by the EU (FEDERAV0013166 and FEDER AV0027831).

## Data

The datasets analysed in the current study are available on request from the corresponding author. It will be shared in open repository when the article will be accepted for publication by a scientific journal.

## Acknowledgments

We extend our thanks to the field workers who helped to collect the data and to the farmers in the Saint-Nectaire PDO production area for allowing access to their plots. We kindly thank Magne Neby, for his constructive advice which allowed us to significantly improve the theoretical framework, specify the materials and methods and the discussion of the results.

## Authors participation

AP, HL, CJ, GSM, ER and VL conceived and designed the experiments. HL, AP, AF, GSM, CJ, MB, ER carried out the field work. HL and AP analysed the data. HL and AP wrote the manuscript and MB and VL provided editorial advice.

## Appendix 1 Argaly labs protocol for DNA extraction & metabarcoding from biological samples

### DNA extraction

#### Soil

DNA from the 43 soil samples was extracted using the phosphate buffer method (Taberlet et al. 2012) with the NucleoSpin® Soil extraction kit (Macherey Nagel GmbH & Co) in a laboratory dedicated to the analysis of environmental samples.

#### Stomach contents

DNA from 37 stomach contents samples was extracted using the NucleoSpin using the NucleoSpin® Soil extraction kit extraction kit (Macherey Nagel GmbH & Co) in a laboratory dedicated for the analysis of environmental samples.

#### Plants

(for the local plant reference database): DNA from 33 plant samples was extracted using the NucleoSpin using the NucleoSpin® Plant II kit (Macherey Nagel GmbH & Co).

All extracted DNA was eluted in a final volume of 100μL.

### Amplification, purification and sequencing

Each soil and stomach content sample was amplified in four replicates with Sper02 plant primers (psbCL gene; primers modified from Riaz et al. 2011; Taberlet et al. personal communication). Each sample was recognised by an eight-base tag for sequence assignment to samples during bioinformatics analysis. All samples were purified using the MinElute purification kit (Qiagen GmbH) and mixed for the sequencing run (Illumina MiSeq run). Each plant tissue sample was amplified in a single replicate with the Sper02 plant primers and sequenced by Sanger sequencing.

### Quality controls

Various controls were carried out at each stage of the protocol in order to detect any contamination and to allow better interpretation of the results.

- For the soil and stomach content samples: three negative extraction controls, four negative PCR controls and twelve bioinformatics blanks.
- For plant samples: one extraction negative control and one PCR negative control.

Successful amplification and purification were verified by agarose gel electrophoresis (E-Gel Precast; Invitrogen).

### Bioinformatics

To analyse the sequencing data, the OBITools suite of bioinformatics programs (Boyer et al. 2016; https://pythonhosted.org/OBITools/welcome.html) was used. These tools are used to assemble paired sequences, assign sequences to the corresponding amplification replicates, filter out artefactual sequences, dereplicate them and finally make a taxonomic assignment for each molecular taxonomic unit(MOTU). ROBITools (https://git.metabarcoding.org/obitools/ROBITools) were then used to examine the results obtained, aggregate the amplification replicates per sample and eliminate contaminants.

For taxonomic assignments, the following steps were carried out:

- The sequencing results were compared separately with the local plant reference database and with the GenBank reference database.
- The assignment ranks obtained were compared in both cases (species, genus, family, order, etc.).
- If these ranks were identical, the assignment from the local database was retained; if these ranks were different, the most precise assignment was retained except when the assignment from the local database was already precise (at least to genus), in which case it was retained so as not to force the assignment of sequences to species that do not exist locally.

## Appendix 2 Overview of plant communities recorded in the different samplings

Data are presented as percentage of plants family and genus or specie (if data available) in studied plots for each sampling type.

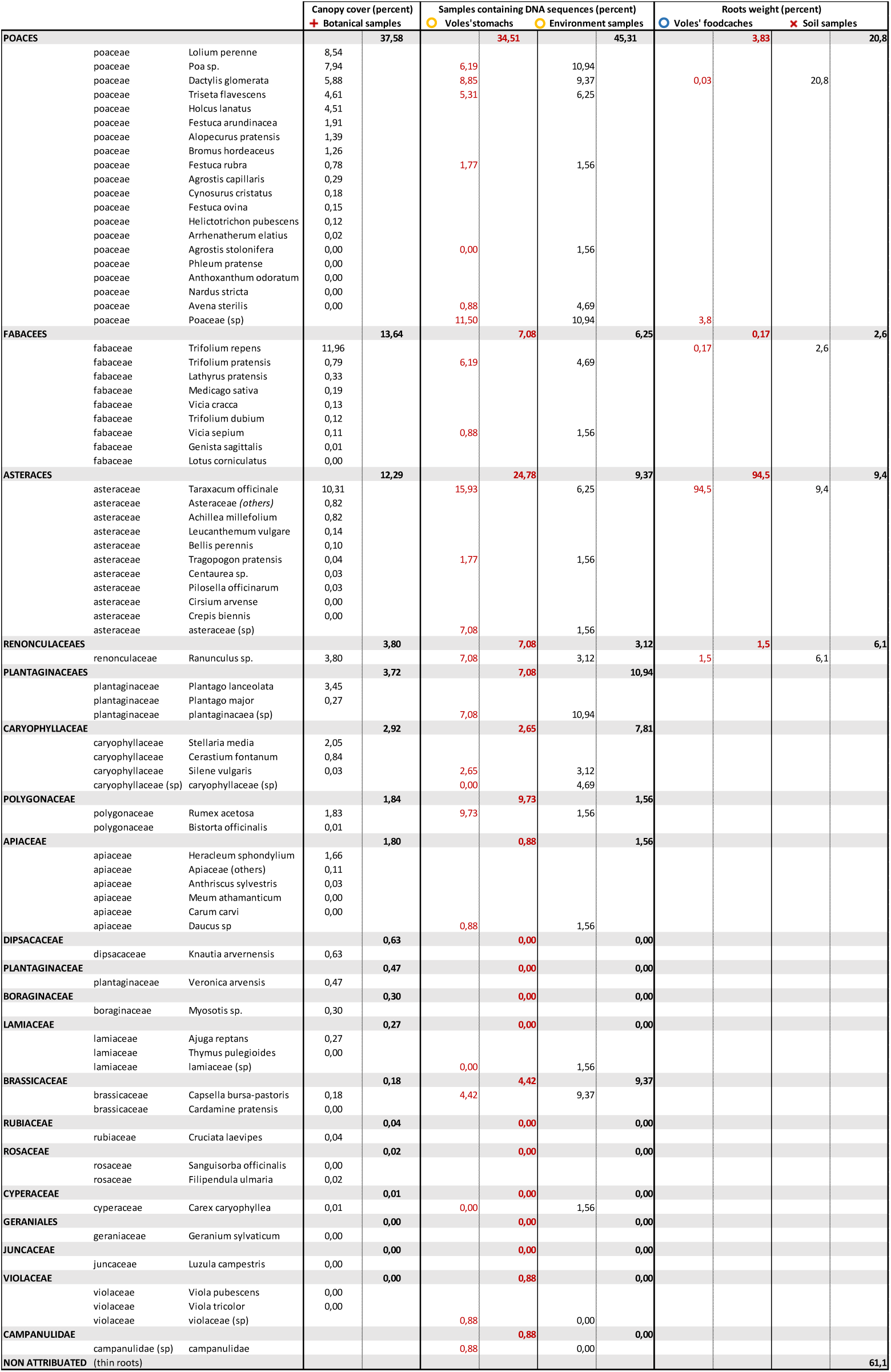

## Notes

### Competing Interest Statement

The authors have declared no competing interest.

### Summary of Updates

The order of the list of authors was wrong in the first version.

